# The representation of two-body shapes in the human visual cortex

**DOI:** 10.1101/637082

**Authors:** Abassi Etienne, Papeo Liuba

## Abstract

Human social nature has shaped visual perception. A signature of the relationship between vision and sociality is a particular visual sensitivity to social entities such as faces and bodies. We asked whether human vision also exhibits a special sensitivity to spatial relations that reliably correlate with social relations. In general, interacting people are more often situated face-to-face than back-to-back. Using functional MRI and behavioral measures in female and male human participants, we show that visual sensitivity to social stimuli extends to images including two bodies facing toward (vs. away from) each other. In particular, the inferior lateral occipital cortex, which is involved in visual-object perception, is organized such that the inferior portion encodes the number of bodies (one *vs.* two) and the superior portion is selectively sensitive to the spatial relation between bodies (facing *vs.* non-facing). Moreover, functionally localized, body-selective visual cortex responded to facing bodies more strongly than identical, but non-facing, bodies. In this area, multivariate pattern analysis revealed an accurate representation of body dyads with sharpening of the representation of single-body postures in facing dyads, which demonstrates an effect of visual context on the perceptual analysis of a body. Finally, the cost of body inversion (upside-down rotation) on body recognition, a behavioral signature of a specialized mechanism for body perception, was larger for facing vs. non-facing dyads. Thus, spatial relations between multiple bodies are encoded in regions for body perception and affect the way in which bodies are processed.

**Public Significance Statement:** Human social nature has shaped visual perception. Here, we show that human vision is not only attuned to socially relevant entities, such as bodies, but also to socially relevant spatial relations between those entities. Body-selective regions of visual cortex respond more strongly to multiple bodies that appear to be interacting (i.e., face-to-face), relative to unrelated bodies, and more accurately represent single body postures in interacting scenarios. Moreover, recognition of facing bodies is particularly susceptible to perturbation by upside-down rotation, indicative of a particular visual sensitivity to the canonical appearance of facing bodies. This encoding of relations between multiple bodies in areas for body-shape recognition suggests that the visual context in which a body is encountered deeply affects its perceptual analysis.

## Introduction

The term *social vision* refers to phenomena in visual perception that mediate human social life (Adams et al., 2010). Among these phenomena, there is a tuning of human vision to certain features or entities that have high social value. Stimuli such as bodies and faces are individuated earlier than other stimuli in infancy (Bonatti et al., 2005). In cluttered environments, they are the most likely to recruit attention (Downing et al., 2004; New et al., 2007). They are also particularly susceptible to the cost of inversion, the detrimental effect on recognition, of seeing a stimulus rotated upside-down (Yin, 1969; Reed et al., 2003). This so-called inversion effect has been linked to an internal representation of the canonical structure of faces or bodies, which makes recognition particularly efficient, and the disruption of such structure (e.g., through inversion) particularly harmful to recognition (Maurer et al., 2002). In the occipito-temporal ventral visual pathway, a number of areas exhibit a preference for faces and bodies, indexed by increased neural activity (Kanwisher et al., 1997; Downing et al., 2001; Gobbini and Haxby, 2007).

Efficient face/body perception, as a building block of social cognition and prerequisite for many social tasks, is an expression of the social function of vision (Adams et al., 2010). Central to social tasks is also the recognition of relations between two or more entities. Third-party relations such as physical interactions must entail rapid discrimination, to activate adaptive behaviors such as defense or assistance/cooperation, and infer group affiliation and conformity (Quadflieg and Koldewyn, 2017). Rapid discrimination might benefit from the fact that some common physical or communicative exchanges reliably correlate with spatial relations: for example, physically interacting people are often close and facing toward, rather than away from, each other.

Behavioral research suggests that key properties of social interaction, such as action coherence and agent/patient-role information, correlate with visuo-spatial features that are accessed quickly in visual events (Hafri et al., 2013, 2018; Glanemann et al., 2016). Moreover, multiple bodies in configurations that imply interaction (e.g., face-to-face), are detected, recognized, and remembered better than unrelated bodies (Ding et al., 2017; Papeo and Abassi, 2019; Papeo et al., 2019; Vestner et al., 2019).

The fact that seemingly interacting bodies are appraised quickly and processed efficiently inspired our hypothesis that visual areas would exhibit increased sensitivity to socially relevant multi-person configurations. Moreover, following evidence for the effects of spatial relations between multiple objects on activations in object-selective visual cortex (Roberts and Humphreys, 2010; Kim and Biederman, 2011; Baeck et al., 2013), we hypothesized that relations between multiple bodies would specifically affect activity in face- and body-selective visual cortex. Previous research has shown that body-selective areas do not distinguish between interacting and non-interacting people based on contextual cues (e.g., accessories or clothing) or action knowledge (i.e., body postures; Quadflieg et al., 2015). Therefore, we exclusively considered a basic visuo-spatial cue of interaction: face-to-face positioning, which is common to many instances of physical and communicative exchanges, and elemental to components of social behavior such as gaze following and shared attention (Baron-Cohen et al., 1997; Birmingham et al., 2009; Graziano and Kastner, 2011).

For face-to-face (facing) and back-to-back (nonfacing) body dyads, we measured two classical indexes used to show visual sensitivity to single bodies: 1) the neural response in body-selective cortex (fMRI experiment); and 2) the magnitude of the inversion effect in body recognition (behavioral experiment). Images of single bodies, facing body dyads, and nonfacing body dyads were presented during fMRI. The same stimuli, upright or inverted upside-down, were also shown during a behavioral experiment involving a visual-categorization task. We reasoned that higher visual sensitivity to facing *vs.* nonfacing dyads would yield stronger neural responses and more accurate representations in body-selective visual cortex, and a larger inversion effect on visual recognition. Neural and behavioral results converged to demonstrate that human visual perception is tuned to two-body configurations in which their relative spatial position suggests an ongoing social exchange.

## Materials and Methods

### fMRI experiment

#### Participants

Twenty healthy adults (12 female; mean age 24.1 years, *SD* = 3.1) took part in the fMRI study as paid volunteers. All had normal or corrected-to-normal vision and reported no history of psychiatric or neurological disorders, or use of psychoactive medications. They were screened for contraindications to fMRI and gave informed consent before participation. The local ethics committee (CPP Sud Est V, CHU de Grenoble) approved the study.

#### Stimuli

Stimuli were grayscale renderings of one or two human bodies in various biomechanically possible poses, seen in a profile view. Poses with fully outstretched limbs were avoided, so that the two halves of each figure contained a roughly comparable number of pixels. Stimuli were created and edited with Daz3D (Daz Productions, Salt Lake City) and the Image Processing Toolbox of MATLAB (MathWorks). Eight unique body poses (fig. 1a), as well as a horizontally flipped version of each pose, formed the single-body set. The center of each image featuring a single body was defined as the mid-point between the most extreme point on the left and the most extreme point on the right along the horizontal axis. Eight facing dyads were created from the eight body poses; therefore, each body was used twice, each time paired with a different second body. The final set of facing dyads included the eight dyads and their horizontally flipped version. Nonfacing dyads were created by swapping the position of the two bodies in each facing dyad (i.e., the body on the left side was moved to the right side and *vice versa*). The mean distance between the two bodies in a dyad was matched across facing and nonfacing stimuli. The centers of the two minimal bounding boxes that contained each figure of a dyad were placed at an equal distance from the center of the image. In addition, we ensured that the distance between the closest points (extremities) of the two bodies was matched across facing and nonfacing dyads (mean_facing_ = 133.25 pixels ±28.68 *SD*; mean_nonfacing_ = 132.25 ±30.03 *SD*; Mann-Whitney *U* = 35.5, *Z* = 0.36, *p* = 0.721, two-tailed). In summary, since the very same bodies were presented, with matched distances, in facing and nonfacing dyads, the relative spatial positioning of bodies (and body parts) was the only visual feature that differed between the two conditions. The final stimulus set included a total of 48 stimuli (16 single bodies,16 facing dyads, and 16 nonfacing dyads). Inside the scanner, stimuli were back-projected onto a screen by a liquid crystal projector (frame rate: 60 Hz; screen resolution: 1024 × 768 pixels, screen size: 40 × 30 cm). For both body dyads and single bodies, the center of the image overlapped with the center of the screen, where a fixation cross was shown. Participants viewed the stimuli binocularly (∼7° of visual angle) through a mirror above the head coil. Stimulus presentation, response collection and synchronization with the scanner were controlled with the Psychtoolbox (Brainard, 1997) through MATLAB (MathWorks).

**Figure 1.**
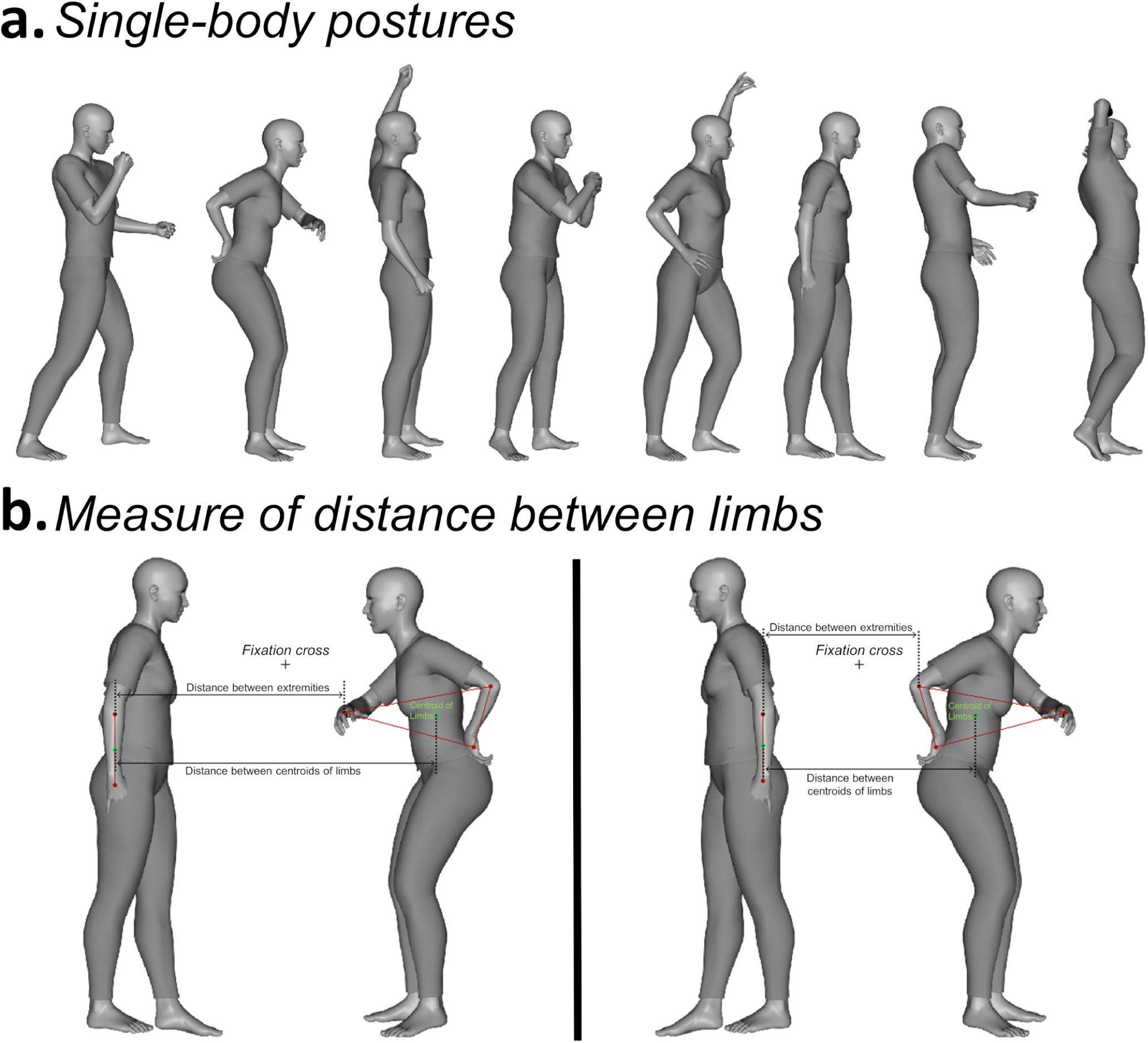
Single-body postures used in the study and coordinates for measuring the distance between limbs. **(a)** The eight body postures presented in the single-body condition and used to create the facing and non-facing dyads. **(b)** Example of coordinates on the two bodies of a dyad, which were considered to quantify the distance between limbs. Distance corresponded to the number of pixels along the *x*-axis, between the centroids of the upper limbs of the two bodies, and between the two closest points on the limbs of the two bodies. The green dots denote the centroids of the hand-to-elbow area delimitated by red dots. The red dot closest to the center of the screen was used to quantify the distance between extremities.

#### Design of the main fMRI experiment

The experiment consisted of two parts: the main fMRI experiment, and a functional localizer task, which we describe below. In the main experiment, facing and nonfacing dyads were presented over three runs, each lasting 7.42 min. Each run consisted of two sequences of 16 blocks (8 facing and 8 nonfacing), for a total of 32 blocks of 6.5 s each. Blocks in the first sequence were presented in a random order, and blocks in the second sequence were presented in the counterbalanced (i.e., reversed) order relative to the first sequence. Thus, blocks that were presented at the end of the first sequence were shown at the beginning of the second sequence, and *vice versa*. Each block featured five repetitions of the same image, randomly alternating between the original view and its flipped version. In each of the three runs, there were therefore two blocks for each dyad, for a total of six blocks for each dyad across the whole experiment. Three additional runs, each lasting 3.93 min, featured single-body images. Each run included 16 blocks (6.5 s each). In each block, the same single-body stimulus was presented five times, alternating between the original view and its flipped version. There were two blocks for each single-body stimulus in each run, and six blocks across the whole experiment. Runs with dyads and runs with single bodies were presented in pseudorandom order to avoid the occurrence of more than two consecutive runs of the same stimulus group. Each run began with a warm-up block (10 s) and ended with a cool-down block (16 s), during which a central fixation cross was presented. Within a run, the onset time of each block was jittered (range of inter-block interval duration: 4.4-13.2 s; total inter-block time for runs with dyads: 211.2 s, for runs with single bodies: 105.6 s) to remove the overlap from the estimate of the hemodynamic response (Dale, 1999). Jittering was optimized using the optseq tool of Freesurfer (Fischl, 2012). During each block, a black cross was always present in the center of the screen, while stimuli appeared for 400 ms, separated by an interval of 900 ms. In a subset (37.5.%) of stimulus and fixation blocks, the cross changed color (from black to red). Participants were instructed to fixate the cross throughout the experiment, and detect and report the color change by pressing a button with their right index finger. This task was used to minimize eye movements and maintain vigilance in the scanner. In total, the main experiment including three runs of dyads and three runs of single bodies and lasted 34.05 min total.

#### Functional Localizer task

In addition to the six experimental runs, participants completed a functional localizer task, with stimuli and a task adapted from the fLoc package (Stigliani et al., 2015) and used in several previous studies (e.g., Natu et al., 2016; Weiner et al., 2017; Gomez et al., 2018). During this task, participants saw 144 grayscale photographs of the following five object classes: 1) body-stimuli including whole bodies (headless female and male bodies in various views and poses) and body parts (hands, arms, elbows, feet, legs and knees); 2) faces (uncropped pictures of female and male adults and children in various views); 3) places (houses and corridors); 4) inanimate objects (various exemplars of cars and musical instruments). Stimuli were presented over three runs (4.47 min each). Each run began with a warm-up block (12 s) and ended with a cool-down block (16 s), and included 60 blocks of four seconds each: 12 blocks for each object class (bodies, faces, places and objects) with eight images per block (500 ms per image without interruption), randomly interleaved with 12 baseline blocks featuring an empty screen. To minimize low-level differences across categories, the view, size, and retinal position of the images varied across trials, and each item was overlaid on a 10.5° phase-scrambled background generated from another image of the set, randomly selected. During blocks containing images, scrambled objects were presented as catch trials, and participants were required to press a button when a scrambled object appeared.

#### Data acquisition

Imaging was conducted on a MAGNETOM Prisma 3T scanner (Siemens Healthcare). T2*-weighted functional volumes were acquired using a gradient-echo echo-planar imaging sequence (GRE-EPI) (repetition time = 2.2 s, echo time = 30 ms, 40 slices, slice thickness = 3 mm, no gap, field-of-view = 220 mm, matrix size = 74 × 74, acceleration factor of 2 with GRAPPA reconstruction and phase encoding set to anterior/posterior direction). For the main experiment and the functional localizer session, we acquired nine runs for a total of 1275 frames per participant. Acquisition of high-resolution T1-weighted anatomical images was performed after the third functional run of the main experiment and lasted 8 min (MPRAGE; 0.8 mm isotropic voxel size, repetition time = 3 s, echo time = 3.7 ms, TI = 1.1 s, field-of-view = 256 × 224 mm, acceleration factor of 2 with GRAPPA reconstruction).

#### Preprocessing

Functional images were preprocessed and analyzed using SPM12 (Friston et al., 2007) in combination with MATLAB and the CoSMoMVPA toolbox (Oosterhof et al., 2016). The first four volumes of each run were discarded, taking into account initial scanner gradient stabilization (Soares et al., 2016). Preprocessing of the remaining volumes involved slice time correction, then spatial realignment and motion correction using the first volume of each run as reference. The maximum displacement was 0.78 mm on the *x*-axis (mean_max_ = 0.27, *SD* = 0.17); 2.54 mm on the *y*-axis (mean_max_ = 0.69, *SD* = 0.67); and 4.60 mm on the *z*-axis (mean_max_ = 1.17, *SD* = 1.04), which did not exceed the Gaussian kernel of 5 mm FWHM used for the spatial smoothing before estimating the realignment parameters (Friston et al., 1996). Final steps included removing low-frequency drifts with a temporal high-pass filter (cutoff 128 s) and spatial smoothing with a Gaussian kernel of 8 mm FWHM for univariate analysis, and of 2 mm FWHM for multivariate analysis. Anatomical volumes were co-registered to the mean functional image, segmented into gray matter, white matter and cerebrospinal fluid in native space, and aligned to the probability maps in the Montreal Neurological Institute (MNI) as included in SPM12. The DARTEL method (Ashburner, 2007) was used to create a flow field for each subject and an inter-subject template, which was registered in the MNI space and used for normalization of functional images.

### Univariate (activity-based) analyses

#### Whole-brain analysis

The blood-oxygen-level-dependent (BOLD) signal of each voxel in each participant was estimated in two random-effects general linear model (RFX GLM) analyses, each including two regressors for the experimental conditions (either single-bodies and dyads, or facing and non-facing dyads), one regressor for fixation blocks, and six regressors for movement correction parameters as nuisance covariates. In one analysis, the effect of the number of bodies presented was assessed with the RFX GLM contrast dyads > single-bodies. In the second analysis, the effect of the spatial relation between bodies in a dyad was assessed with the RFX GLM contrasts facing > nonfacing dyads and nonfacing > facing dyads. The statistical significance of second-level (group) effects was determined using a voxelwise threshold of *p* ≤ 0.001, family-wise error corrected at the cluster level.

#### Voxel-by-voxel conjunction analysis in Lateral Occipital Cortex (LOC)

Following up on the effect of spatial relations between objects specifically found in the LOC (Baeck et al., 2013; Roberts & Humphreys, 2010), we carried out a voxel-by-voxel conjunction analysis (Friston et al., 2005) across this area in order to disentangle the effects of body number and spatial relation, and to identify the aspects of LOC that showed an effect of spatial relation. The left and right functional maps from the group-level contrasts dyads > single-bodies and facing dyads > nonfacing dyads were superimposed on the anatomical masks of the left and right LOC, respectively. Within the left and right mask, three functional maps were defined, which included voxels with a significant level of activation (threshold: *p* = 0.001) only in the facing > nonfacing contrast, only in the dyads > single-bodies contrast, or in both contrasts (Friston et al., 2005).

#### Definition of Regions of interest (ROIs) and ROIs analysis

We asked whether spatial relations between bodies in a dyad are encoded in the same neural structures which are specialized for body perception, independently identified at the individual subject level. To this end, ROIs were defined by entering the individual subjects’ data, registered during the functional localizer task, into a GLM with four regressors for the four object class conditions (bodies, faces, places and objects), one regressor for baseline blocks, and six regressors for movement correction parameters as nuisance covariates. Three bilateral masks of the inferior LOC, the temporal occipital fusiform cortex and the inferior parahippocampal cortex were created using FSLeyes (McCarthy, 2019) and the Harvard-Oxford Atlas (Desikan et al., 2006) through FSL (Jenkinson et al., 2012). Within each mask of each participant, we selected the voxels with significant activity (threshold: *p* = 0.05) for the contrasts of interest. In the LOC, we identified the extrastriate body areas (EBA) with the contrast of bodies > objects and the occipital face area (OFA) with the contrast of faces > objects. In the temporal occipital fusiform cortex, we identified the fusiform face area (FFA) with the contrast of faces > objects and the fusiform body area (FBA) with the contrast of body > objects. In the parahippocampal cortex, the parahippocampal place area (PPA) was localized with the contrast of places > objects. The PPA has been shown to discriminate between object sets based on internal spatial relations, with particularly strong responses to sets that form familiar scenes (Aminoff et al., 2007; Kaiser et al., 2014). We included this ROI to determine if it serves a general function in processing spatial relations between multiple objects or, rather, if spatial relations between bodies are encoded selectively in body-selective visual cortical areas. All of the voxels in a mask and in its contralateral homologue that passed the threshold were ranked by activation level based on *t* values. The final ROIs included up to 200 (right or left) best voxels (mean number of voxels in the EBA and PPA: 200; in the FBA: 191.2 ± 39.35 *SD*; in the OFA: 198.15 ± 8.27 *SD;* in the FFA: 198.4 ± 7.16 *SD*). Finally, a bilateral mask of the early visual cortex (EVC) was created using a probabilistic map of visual topography in human cortex (Wang et al., 2015). After transforming the mask in each subject space, the 200 right or left voxels with the highest probability were selected to form the EVC-ROI. From the six ROIs of each participant, the mean neural activity values (mean β-weights minus baseline) for facing and nonfacing dyads were extracted and analyzed in a 2 spatial relation x 6 ROI repeated-measures ANOVA, run with Statistica (StatSoft Europe, Hamburg). To address the lateralization of the effects, a second repeated-measures ANOVA included, in addition to the above factors, the hemisphere (left vs. right). For this analysis, the mean β-weights were extracted separately from the left and right ROIs, each consisting of a maximum of 200 voxels with significant activity (threshold: p = 0.05) for the contrast of interest, or with the highest probability in the maps of visual topography for the EVC. When a functional ROI could not be identified (the left OFA for two participants, the left FBA for one participant and the right FBA for one participant), individual β-weights were extracted from a mask corresponding to the group-level activation for the contrast of interest.

### Multivariate pattern analyses (MVPA)

#### Multi-class classification of single bodies from body dyads

Using MVPA, we examined body representation as a function of the spatial relation between two bodies. In particular, in the six ROIs, we measured how well single body postures could be discriminated in facing *vs.* nonfacing dyads. ROIs were defined as above, using the data from the functional localizer task. Spatial smoothing of 2 mm FWHM was applied, to be consistent with the 2 mm FWHM preprocessing used for MVPA (mean number of voxels in EBA: 198 ± 2.85 *SD*; in FBA: 196 ± 9.22 *SD*; in OFA: 198.3 ± 2.79 *SD*; in FFA: 198.45 ± 2.35 *SD*; in PPA: 193.3 ± 18.01 *SD*; in EVC: 188.1 ± 10.94 *SD*). For each participant, in each ROI, we estimated 48 multivariate β-patterns for facing dyads and 48 for nonfacing dyads (96 patterns in total for three runs), along with 48 β-patterns for single bodies (for three runs). β-patterns were normalized run-wise to avoid spurious correlations within runs (Lee and Kable, 2018). A support vector machine (SVM) classifier (LIBSVM, Chang and Lin, 2011) implemented in the CosmoMVPA toolbox (Oosterhof et al., 2016) was trained to classify the patterns of eight classes corresponding to the eight single bodies, using six samples per class. It was then tested on the classification of the 48 patterns corresponding to facing dyads, using a one-against-one approach and voting strategy, as implemented in LIBSVM. In 48 testing iterations, a pattern corresponding to a facing dyad was classified in one of the eight classes of single bodies. For each participant, classification accuracy values were averaged across all iterations. Since the test items each included two bodies, classification of each body could be correct in two out of eight cases; therefore, the chance level was 25%. This analysis was repeated using the patterns for nonfacing dyads as the test set. For each ROI, individual classification accuracy values for train-and-test on facing dyads and for train-and-test on nonfacing dyads were tested against chance with *t* tests (α = 0.05, two-tailed).

#### Multi-class classification of synthetic-mean patterns from real patterns

Using MVPA, we tested the discrimination of individual body dyads in body-selective cortical regions and the quantitative relationship between the representation of each body dyad and the representation of the two bodies forming the dyad. In particular, we assessed whether neural patterns for facing and nonfacing dyads could be modeled as the mean of the neural patterns measured for each of the two constituent bodies, or if this linear relationship changed as an effect of the spatial relation between bodies (see Kubilius et al., 2015). To do so, for each ROI, for each presentation of a dyad, we created a synthetic pattern by averaging the β-patterns associated with the two single bodies in the dyad. An SVM classifier (LIBSVM, Chang and Lin, 2011) was trained with the 48 synthetic β-patterns to classify eight unique class of patterns (corresponding to the eight unique pairings of bodies), and tested on the 48 β-patterns actually measured for facing dyads (six presentations for each of the unique dyads) and, separately, on the 48 β-patterns actually measured for nonfacing dyads. For each test, the chance level was 12.5 % (one out of eight patterns). For each ROI, individual classification accuracy values for train-and-test on facing dyads and for train-and-test on nonfacing dyads were tested against chance using *t* tests (α = 0.05, two-tailed).

### Visual recognition experiment

#### Participants

This experiment involved 16 participants of the fMRI-experiment sample, who volunteered to return to the lab, in addition to five new participants, for a total of 21 participants (15 females; mean age 24.10 years ± 2.96 *SD*). The sample size was decided based of the effect size *η*_*p*_^*2*^ =0.453 (*β* = 0.80, alpha = 0.05; G*Power 3.1, Faul et al., 2007) obtained in a previous study that used similar paradigm and design (Papeo et al., 2017).

#### Stimuli

In addition to the 48 images of body-stimuli (16 single bodies, 16 facing dyads and 16 non-facing dyads) used in the fMRI experiment, the current experiment included 48 images of single chairs or pairs of chairs. Chair stimuli were created from eight grayscale exemplars of chairs and their horizontally flipped versions, which were combined in 16 pairs of facing, and 16 pairs of non-facing chairs. Body and chair stimuli were inverted upside down, yielding a total of 192 stimuli presented against a grey background. An equal number of masking stimuli was created, consisting of high-contrast Mondrian arrays (11° × 10°) of grayscale circles (diameter 0.4°-1.8°).

#### Design and Procedures

The task included two identical runs, each containing 32 trials for each of the twelve conditions (upright and inverted single, facing and nonfacing bodies and chairs), presented in a random order. Each stimulus appeared twice in a run. Participants sat on a height-adjustable chair, 60 cm away from a computer screen, with their eyes aligned to the center of the screen (17-in. CRT monitor; 1024 × 768 pixel resolution; 85-Hz refresh rate). Stimuli on the screen did not exceed 7° of visual angle. Each trial included the following sequence of events: blank screen (200 ms), fixation cross (500 ms), blank screen (200 ms), target stimulus (30 ms), mask (250 ms) and a final blank screen that remained until the participant gave a response. The next trial began after a variable interval between 500 and 1000 ms. For each trial, participants responded by pressing one of two keys on a keyboard in front of them (“1” with the index finger for “bodies”, or “2” with the middle finger for “chair”; this stimulus-response mapping was counterbalanced across participants). Every 32 trials participants were invited to take a break. Two blocks of familiarization trials preceded the experiment. In the first block, four stimuli per condition were shown for 250 ms, so that the participant could see the stimuli clearly. In the second block, eight stimuli per condition were shown for 30 ms, as in the actual experiment. The instructions for the familiarization blocks were identical to those of the actual experiment. The experiment lasted ∼40 min. Stimulus presentation and response collection (accuracy and RTs) were controlled with Psychtoolbox (Brainard, 1997) through MATLAB.

#### Behavioral data analyses

Data from one participant were discarded because the average RT was more than 2.5 *SD* above the group mean. Mean accuracy (mean proportion of correct responses) and RTs for the remaining 20 participants were used for the analysis of the body inversion effect, in a 3 stimulus x 2 orientation repeated-measures ANOVA. Comparisons between critical conditions were performed with pairwise *t* tests (α = 0.05, two-tailed). The same analysis was repeated on RTs, after removing trials in which the participant’s response was inaccurate or 2 *SD* away from the individual mean (9.6% of the total number of trials).

#### Correlation between behavioral and neural effects of facing (vs. non-facing) dyads

For the 15 subjects who participated in both the behavioral and the fMRI session of the study, we computed the correlation between the difference in the inversion effect for facing dyads *vs*. nonfacing dyads [(accuracy for upright minus inverted facing dyads) - (accuracy for upright minus inverted nonfacing dyads)], and the difference in the activation for facing *vs.* nonfacing dyads in the body-selective ROIs (beta weight for facing minus beta weight for nonfacing dyads). The Pearson correlation coefficients computed for all participants were Fisher-transformed and entered in a one-sample *t* test against 0 (two-tailed).

## Results

### fMRI

#### Effects of number and spatial relation across the whole brain

We first examined the brain regions responsive to the number of bodies and to the spatial relation in body dyads by comparing dyads vs. single bodies and facing vs. nonfacing dyads, respectively (Table 1). A large bilateral cluster, centered in the inferior temporal gyrus, responded to dyads more strongly than to single bodies. This activation spread more posteriorly into the lateral occipital cortex. A stronger response to facing than nonfacing dyads was found in a bilateral cluster encompassing the posterior middle temporal gyrus and the middle occipital gyrus. In addition, relative to nonfacing dyads, facing dyads elicited stronger bilateral prefrontal activity, centered in the anterior middle frontal gyrus (Fig. 2a). No significant increase of activity for nonfacing dyads compared with facing dyads was found, implying that no region was activated more, or selectively, for nonfacing dyads.

**Table 1.** Activations for the whole-brain contrasts. Location and significance of clusters showing stronger responses to dyads relative to single bodies, and to facing dyads relative to nonfacing dyads.

| Contrasts | Hemispheres | Locations | Peak coordinates |  |  | *t* | Cluster-Wise<br>FWE | Cluster<br>size |
| --- | --- | --- | --- | --- | --- | --- | --- | --- |
|  |  |  | x | y | z |  |  |  |
| Dyads ><br>Individuals | Right | Inferior Temporal Gyrus | 47 | -71 | -3 | 7.82 | <0.001 | 3232 |
|  | Left | Middle Temporal Gyrus | -51 | -74 | 8 | 6.21 | <0.001 | 2348 |
| Facing ><br>Non-facing | Right | Middle Frontal Gyrus | -41 | 44 | -5 | 7.59 | <0.001 | 3307 |
|  |  | Middle Occipital Gyrus | -41 | -72 | 2 | 4.75 | <0.001 | 1335 |
|  | Left | Middle Frontal Gyrus | 45 | 33 | 24 | 4.87 | 0.004 | 805 |
|  |  | Middle Temporal Gyrus | 48 | -59 | 6 | 5.11 | 0.011 | 652 |

Note. Results are cluster-wise corrected using FWE. Peak coordinates are in MNI space.

**Figure 2.**
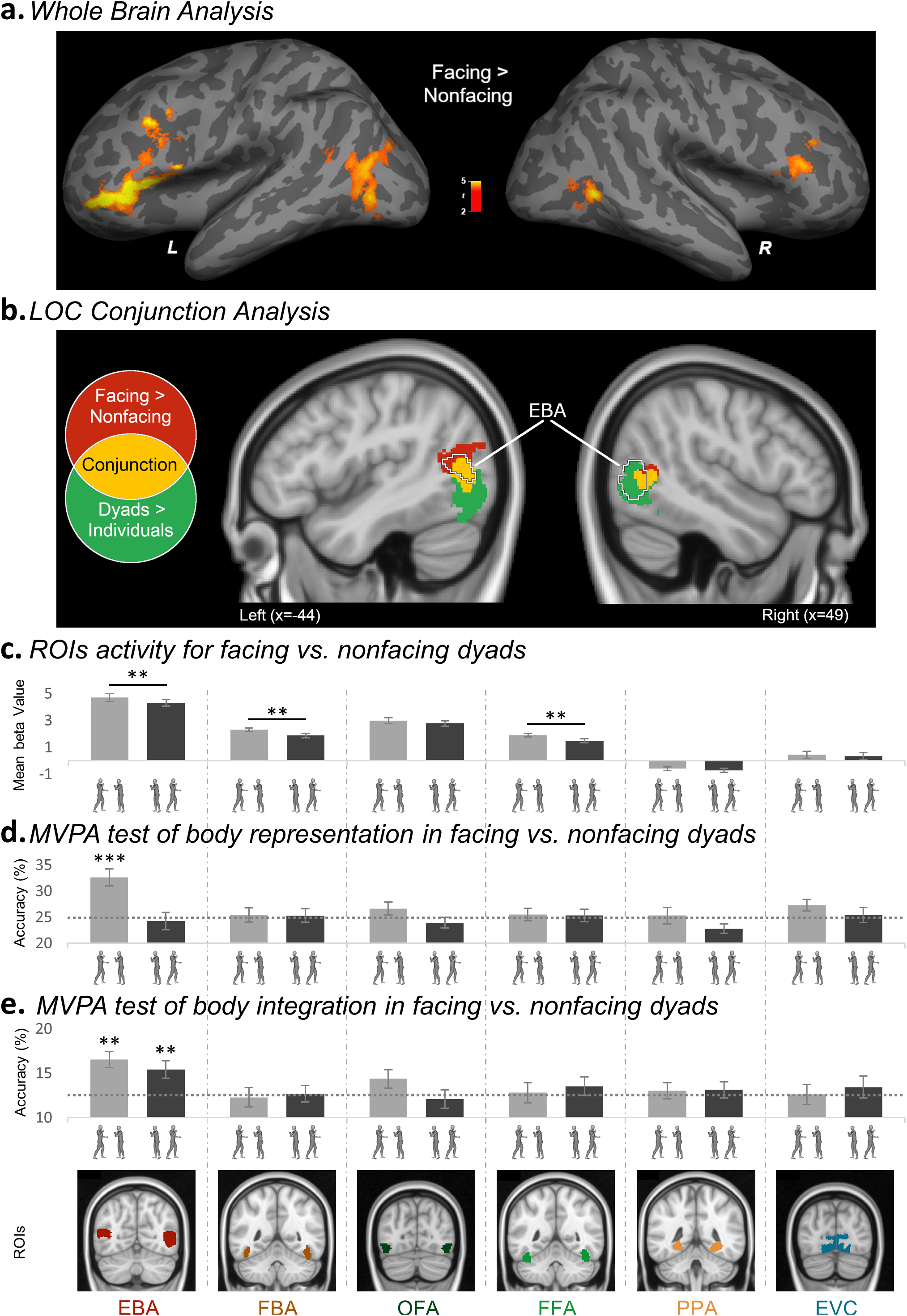
Increased activity for facing *vs.* non-facing dyads in the whole-brain and the ROIs, and representational sharpening in the EBA. **(a)** Left (L) and right (R) group random-effect map (N = 20) for the contrast facing dyads > non-facing dyads. The color bar indicates *t* values. **(b)** Conjunction of the group random-effect maps of the contrasts facing dyads > non-facing dyads (effect of spatial relation) and dyads > single-bodies (effect of number), in the bilateral LOC. Highlighted in red are the voxels showing significant effect of spatial relation only; highlighted in yellow are the voxels showing the effect of both spatial relation and number (conjunction); highlighted in green are the voxels showing the effect of number only. The EBA location corresponds to the group random-effect contrast bodies > objects using the data of the functional localizer task. (**c)** Average beta values (± within-subjects normalized *SEM*) across participants, in each individually defined ROI (EBA, FBA, OFA, FFA, PPA and EVC), in response to facing and non-facing dyads. \*\**p* ≤ 0.01. **(d)** Classification accuracies (± within-subjects normalized *SEM*) for multi-class cross-decoding of single bodies patterns from facing dyads and non-facing dyads (training on patterns of single-bodies, test on patterns of facing dyads and of non-facing dyads), in each ROI. The horizontal dashed grey bar represents the chance level (25%). **(e)** Classification accuracies (± within-subjects normalized *SEM*) for multi-class cross-decoding of synthetic-mean patterns (mean of patterns for the two single bodies) from facing dyads and non-facing dyads (training on the synthetic-mean patterns, test on patterns of facing dyads and non-facing dyads), in each ROI. Horizontal dashed bars represent the chance level (12.5%). Asterisks indicate accuracy of classification significantly above chance: \*\**p* ≤ 0.01; \*\*\**p* ≤ 0.001. For illustration purposes, ROIs correspond to the results of the group-level random-effect contrasts of bodies > objects for the EBA and the FBA, faces > objects for the OFA and the FFA, and places > objects for the PPA and the probabilistic map of EVC.

#### Effects of number and spatial relation across the LOC

The group-level contrasts revealed widespread effects of both number (single body vs. dyad) and spatial relation (facing vs. non-facing dyads) in the LOC. This observation confirmed the functional relevance of the LOC in processing multiple-body configurations, which we hypothesized after previous reports of the effects of object positioning in this region (Roberts and Humphreys, 2010; Kim and Biederman, 2011; Baeck et al., 2013). With the current voxel-by-voxel conjunction analysis (Friston et al., 2005), we sought to disentangle the effect of body number from the effect of spatial relation, to highlight the aspects of this region that showed an effect of spatial relation. The conjunction of the statistical maps of the whole-brain contrasts dyads > individuals and facing > nonfacing dyads, superimposed to the anatomically-defined bilateral LOC, revealed three consecutive regions along the inferior-to-superior axis, with three different response profiles (Fig. 2b). Voxels in the inferior part of LOC showed an effect of number, with a stronger response to dyads than single bodies. The middle region included voxels that responded more strongly to dyads than single bodies, and to facing dyads than nonfacing dyads. Finally, the superior region showed a selective effect of spatial relation, responding to facing dyads more strongly than to nonfacing dyads. As illustrated in Fig. 2b, the middle and superior regions, which showed sensitivity to spatial relation, overlapped with the body-selective EBA, as independently identified in the following ROI analysis. Thus, besides addressing the anatomical and functional relationship between the effects of number and spatial relation, the voxel-by-voxel analysis provided the first indication that spatial relations between bodies were represented specifically in the body-selective aspect of the LOC.

#### Effects of the spatial relation in the face- and body-selective cortex

Stronger activity for facing *vs.* nonfacing dyads was found in the EBA, the body-selective cortex within the LOC. To further investigate this finding and overcome the limits of group analysis in the definition of anatomical-functional correspondences (Saxe et al., 2006), we examined the effect of the spatial relation between bodies in the functionally defined face/body-selective cortex in the EBA, FBA, FFA and OFA. In addition, we tested the functionally defined PPA, a region that shows sensitivity to spatial relations within object sets (Aminoff et al., 2007; Kaiser et al., 2014), and the EVC, as defined by a probabilistic map of visual topography (Wang et al., 2015).

As shown in Fig. 2c, the relative positioning of bodies in a dyad affected the neural response in the EBA, FBA and FFA, but not in the OFA. Confirming this observation, the 2 spatial relation by 6 ROI repeated-measures ANOVA revealed main effects of spatial relation, *F*(1,19) = 6.05, *p* = 0.024, *η*_p_^2^ = 0.24, and of ROI, *F*(5,95) = 71.59, *p* < 0.001, *η*_p_^2^ = 0.79, which were qualified by a significant two-way interaction, *F*(5,95) = 5.24, *p* < 0.001, *η*_p_^2^ = 0.22. The interaction reflected a significantly stronger response to facing *vs.* nonfacing dyads in the EBA, *t*(19) = 3.09, *p* = 0.006, in the FBA, *t*(19) = 3.62, p = 0.002, and in the FFA, *t*(19) = 3.60, *p* = 0.002, but not in the OFA, *t*(19) = 1.64, p = 0.117, in the PPA, *t*(19) = 1.51, *p* = 0.148, and in the EVC, *t*(19) < 1, *n.s.*. Thus, the EBA, FBA and FFA responded to facing dyads more strongly than to nonfacing dyads. An ANOVA with hemisphere (left vs. right) as an additional within-subjects factor, showed no effect of hemisphere in the interaction between spatial relation and ROI (effect of spatial relation: *F*(1,19) = 5.04, *p* = 0.037, *η*_p_^2^ = 0.21; effect of ROI: *F*(5,95) = 78.27, *p* < 0.001, *η*_p_^2^ = 0.80; effect of hemisphere: *F*(1,19) = 8.24, *p* = 0.010, *η*_p_^2^ = 0.30; spatial relation by ROI: *F*(5,95) = 8.00, *p* < 0.001, *η*_p_^2^ = 0.30; spatial relation by hemisphere: *F*(1,19) = 1.32, *p* = 0.264, *η*_p_^2^ = 0.07; ROI by hemisphere, *F*(5,95) = 1.57, *p* = 0.175, *η*_p_^2^ = 0.08; spatial relation by ROI by hemisphere, *F*(5,95) = 1.26, *p* = 0.288, *η*_p_^2^ = 0.06).

#### No effect of limb eccentricity on the response to body dyads across ROIs

Isolated limbs can trigger strong responses in the body-selective visual cortex (Taylor et al., 2007). Since participants were instructed to fixate the center of the screen, the amount of limb information available in the participants’ central view could vary from trial to trial. The following analyses were carried out to assess: *a)* whether the distance of the limbs from the central fixation point differed systematically between facing and nonfacing dyads; and *b)* to what extent limb eccentricity could account for the responses to facing and nonfacing dyads in body-selective cortex. We focused on the upper limbs as the central fixation was placed in correspondence with the upper body (i.e., around the chest-area). First, we measured the distance between the upper limbs of the two bodies as a proxy for the amount of limb information in the central area. By our reasoning, the shorter the distance between the limbs of the two bodies on the left and right of fixation, the closer the limbs to the fixation cross; therefore, the higher the amount of limb information in the center of the screen.

Distance was quantified as the number of pixels along the *x*-axis (horizontal distance) both between the upper limbs’ centroids (from hands to elbows) of the two bodies, and between the two closest points on the limbs of the two bodies (extremities) (see Fig. 1b). Statistical analysis (two-tailed Mann-Whitney *U* test) showed that the distance (number of pixels) between upper limbs of the two bodies was comparable between facing and nonfacing dyads, considering both the centroids (mean_facing_ = 307.88 ±59.10 *SD*; mean_nonfacing_ = 324.50 ±57.25; *U* = 28, *Z* = 0.36, *p* = 0.721) and the extremities (mean_facing_ = 218.88 ±62.50; mean_nonacing_ = 258.63 ±64.69; *U* = 22.5, *Z* = 0.98, *p* = 0.328).

While this outcome ruled out a role of limb eccentricity in the increased activation for facing *vs.* nonfacing dyads, we nevertheless assessed to what extent this property of the stimuli could predict activity in body-selective cortex. Therefore, for each participant, in each body-selective ROI, we computed the coefficient of correlation (Pearson’s *rho*) between the limb distance in each presentation of a dyad (96 presentations: 48 facing and 48 nonfacing) and the corresponding brain activity (beta weight). Individual correlation coefficients were Fisher-transformed and entered in a one-sample *t* test (two-tailed). We found no correlation in the EBA (with centroids: *mean rho* = −0.02, *t*(19) < 1, *n.s.*; with extremities: *mean rho* = −0.01, *t*(19) < 1, *n.s.*), or in other face/body-selective ROIs (with centroids in the OFA: *mean rho* = −0.02, *t*(19) = −1.04, *p* = 0.31; all other tests: *t*(19) < 1, *n.s.*).

The role of limb eccentricity on the neural response to dyads was further investigated with an analysis of covariance (ANCOVA): in each ROI, we tested the difference in activity (β-weight) for facing *vs.* nonfacing dyads (categorical factor), while statistically controlling for the effect of distance between limbs (covariate). We found that the effect of positioning remained significant even after removing the effect of the covariate in the EBA (centroids: *F*(1,13) = 9.57, *p* = 0.009, *η*_*p*_^*2*^ = 0.42; extremities: *F*(1,13) = 10.10, *p* = 0.007, *η*_*p*_^*2*^ = 0.44), as well as in the FBA (centroids: *F*(1,13) = 5.80, *p* = 0.032, *η*_*p*_^*2*^ = 0.31; extremities: *F*(1,13) = 5.29, *p* = 0.039, *η*_*p*_^*2*^ = 0.29) and FFA (centroids: *F*(1,13) = 5.74, *p* = 0.032, *η*_*p*_^*2*^ = 0.31; extremities: *F*(1,13) = 5.14, *p* = 0.041, *η*_*p*_^*2*^ = 0.28).

In summary, we found no evidence for an effect of the distance of limbs from the central fixation on the response to dyads in the ROIs that showed an effect of body positioning (the EBA, FBA and FFA). Obviously, the lack of limb eccentricity bias does not exclude the possibility that body-selective areas can encode information about limb/body positioning in the visual field. On the contrary, we argue for the opposite point: the effect of body positioning in the EBA, FBA and FFA means that those areas encode body and/or body-part location *de facto*, in addition to the object category (body *vs.* other; see Schwarzlose et al., 2008).

### Body representation in facing *versus* non-facing dyads

Three analytic strategies (whole-brain analysis, voxel-by-voxel conjunction analysis in LOC, and ROI analysis) converged in showing a particular responsiveness to dyads with facing (seemingly interacting) bodies in the EBA. Next, we sought to shed light on the mechanism underlying this increase of neural activity. Since the EBA is known to encode the specific body shape, or posture, in a percept (Downing et al., 2006; Downing and Peelen, 2011), we began by asking whether this functionality was enhanced in the context of facing dyads. We reasoned that, in a facing dyad, a body could provide a context to the other body, which could enrich (i.e., by disambiguating) the representation of individual postures (see Neri et al., 2006), and/or put particular pressure on their discrimination for the purpose of understanding the ongoing interaction. We addressed this question using MVPA with a multi-class cross-decoding scheme that measured, in each ROI, how well single body postures, used for the training, could be discriminated from facing or nonfacing dyads, used in the test. In each test, a pattern corresponding to a (facing or nonfacing) dyad was classified in one of eight classes of single bodies. We found that single bodies could be discriminated accurately only from facing dyads, and only with neural patterns extracted from the EBA (Fig. 2d). In particular, in the EBA, classification accuracy was significantly above chance for facing dyads (mean accuracy = 32.60%, *t*(19) = 4.69, *p* < 0.001) but not for nonfacing dyads (mean accuracy = 24.27%, *t*(19) < 1, *n.s.*). Classification in the former condition was significantly higher than in the latter one (*t*(19) = 3.19, *p* = 0.005). The same result was obtained with different voxel counts ranging from 50 to 500 (Fig. 3a). The same analysis, in the other ROIs, yielded no significant effects (FBA: test on facing dyads: mean accuracy = 25.42%, *t*(19) < 1, *n.s.*, on nonfacing dyads: mean accuracy = 25.31%, *t*(19) < 1, *n.s.*; OFA: test on facing dyads: mean accuracy = 26.67%, *t*(19) = 1.32, *p* = 0.202, on nonfacing dyads: mean accuracy = 23.96%, *t*(19) < 1, *n.s.*; FFA: test on facing dyads: mean accuracy = 25.52%, *t*(19) < 1, *n.s.*, on nonfacing dyads: mean accuracy = 25.31%, *t*(19) < 1, *n.s.*; PPA: test on facing dyads: mean accuracy = 25.31%, *t*(19) < 1, *n.s.*, on nonfacing dyads: mean accuracy = 22.81%, *t*(19) = −2.02, *p* = 0.058; EVC: test on facing dyads: mean accuracy = 27.29%, *t*(19) = 1.98, *p* = 0.063, on nonfacing dyads: mean accuracy = 25.42%, *t*(19) < 1, *n.s.*).

**Figure 3.**
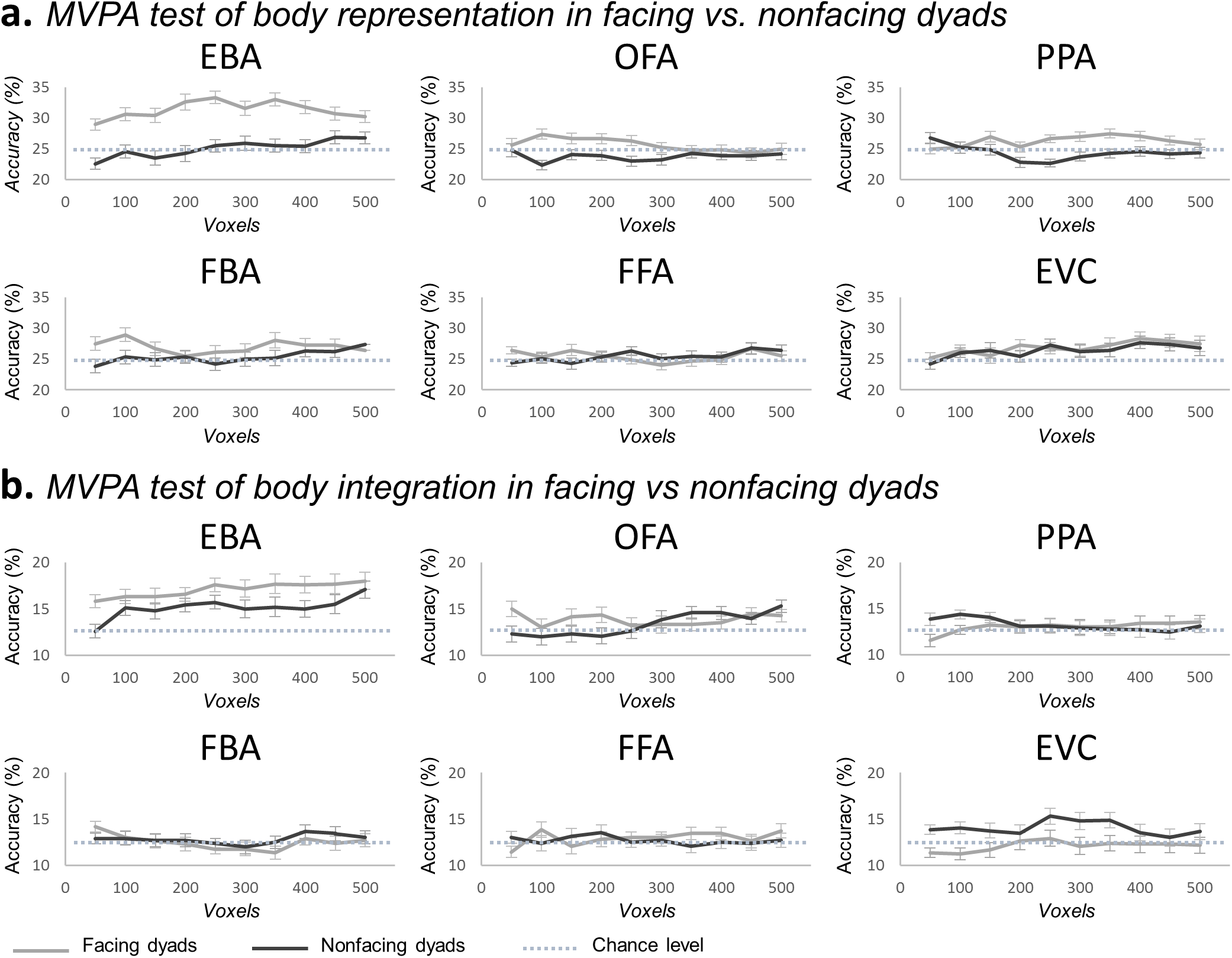
Results of ROI-based MVPA analyses for voxel count from 50 to 500. **(a)** Classification accuracies (± within-subjects normalized *SEM*) for multi-class cross-decoding of single bodies patterns in facing dyads (light grey lines) and non-facing dyads (dark grey lines), in each individually defined ROI (EBA, FBA, OFA FFA and PPA and EVC) using voxels counts from 50 to 500 voxels. Horizontal dashed bars represent the chance level (25%) **(b)** Classification accuracies (± within-subjects normalized *SEM*) for multi-class cross-decoding of synthetic-mean patterns from facing dyads (light grey lines) and non-facing dyads (dark grey lines), in each ROI using voxels counts from 50 to 500 voxels. Horizontal dashed bars represent the chance level (12.5%).

### Body integration in facing *vs.* non-facing dyads

In the light of recent research (Baeck et al., 2013; Kubilius et al., 2015; Brandman and Yovel, 2016; Harry et al., 2016; Baldassano et al., 2017; Kaiser and Peelen, 2018), we tested whether the increased activity for seemingly interacting (*vs.* independent) bodies in the EBA, FBA and FFA, could reflect additional integrative processes. These are expected to cause the representation of a multi-part stimulus (e.g., a dyad) to deviate from the *default* linear combination of information (e.g., mean) from single components of the stimulus (e.g., single bodies). The result of MVPA (Fig. 2e) showed that, exclusively in the EBA, after training on the synthetic-mean β-patterns, classification of real β-patterns was successful for both facing dyads (mean accuracy = 16.56%, *t*(19) = 3.37, *p* = 0.003) and nonfacing dyads (mean accuracy = 15.42%, *t*(19) = 3.07, *p* = 0.006), with no difference between classification accuracy in the two conditions (*t*(19) < 1, *n.s.*). This was consistent across different voxel counts ranging from 50 to 500 (Fig. 3b). The same analysis yielded no significant effects in the other ROIs, including those that showed an effect of spatial relation (the FBA and FFA; all *t*(19) < 1, *n.s.*). Thus, the EBA was the only ROI to show successful encoding of individual body dyads in both the facing and the nonfacing condition. Moreover, in both conditions, information about single bodies was preserved, implying no additional integrative processes for facing dyads. We emphasize that nonfacing dyads were successfully discriminated in the EBA only when the training was based on information about both bodies (in the form of synthetic patterns), but not when it was based on information about one single-body of the dyad. Most likely, this reflects the fact that synthetic patterns carried richer information about the dyads than the patterns associated with one single body only. Finally, we observed that, while they responded to facing dyads more strongly than to nonfacing dyads, the FBA and FFA did not carry enough information about whole body postures to discriminate between different dyads or single bodies.

In a final analysis, we extended the test of integrative processes to the other components of the network triggered by facing dyads, in order to attempt a functional characterization of their activation. We used the MNI group-level peak coordinates (for the whole-brain contrast facing > nonfacing dyads) of the clusters in the middle frontal gyrus (right: −41/44/-5; left: 45/33/24) and in the left middle temporal gyrus (48/-59/6), to create, for each participant, three spherical ROIs (5 mm radius, 171 voxels). In the new ROIs, we performed the above MVPA with cross-decoding of synthetic-mean and actual β-patterns, used for training and test, respectively. Selectively in the right middle frontal gyrus, classification was significantly above chance in the nonfacing-dyad condition (mean accuracy = 15.21%, *t*(19) = 3.07, *p* = 0.006), but not in the facing-dyad condition (mean accuracy = 11.15%, *t*(19) = −1.51, *p* = 0.148), and classification in the nonfacing condition was significantly higher than in the facing condition (*t*(19) = 3.22, *p* = 0.004). In all other ROIs, classification was at chance in both conditions (left middle frontal gyrus: facing dyads: mean accuracy = 11.77%, *t*(19) < 1, *n.s.*; nonfacing dyads: mean accuracy = 11.15%, *t*(19) = −1.55, *p* = 0.137; left middle temporal gyrus: facing dyads: mean accuracy = 13.54%, *t*(19) = 1.04, *p* = 0.309; non-facing dyads: mean accuracy = 12.92%, *t*(19) < 1, *n.s.*).

### Visual recognition

#### The inversion effect for facing versus non-facing body dyads

To show the hypothesized tuning for facing dyads, we examined the inversion effect, a behavioral signature of visual sensitivity. Extensive research has shown that the more the visual system is attuned to the canonical (upright) appearance of a stimulus, the higher the cost of disrupting that canonical configuration (e.g., through inversion), on recognition (Diamond and Carey, 1986; Gauthier et al., 2000). The inversion effect for facing *vs.* nonfacing dyads was measured during recognition of bodies under low-visibility conditions, achieved with short stimulus presentation (30 ms) and backward masking. We found that the magnitude of the inversion effect on both accuracy and RTs varied as a function of the stimulus (Fig. 4a and 4b). A 3 stimulus (single body vs. facing dyads vs. nonfacing dyads) by 2 orientation (upright vs. inverted) repeated-measures ANOVA on accuracy values showed significant effects of stimulus (*F*(2,38) = 5.52, *p* = 0.008, *η*_p_^2^ = 0.22), and orientation (*F*(1,19) = 8.52, *p* = 0.009, *η*_p_^2^ = 0.31), and a significant two-way interaction (*F*(2,38) = 3.94, *p* = 0.028, η_p_^2^ = 0.17). The inversion effect was significant for single bodies (*t*(19) = 2.75, *p* = 0.013), and facing dyads, (*t*(19) = 3.31, *p* = 0.004), but only marginal for nonfacing dyads (*t*(19) = 2.01, *p* = 0.058); importantly, it was significantly larger for facing than for nonfacing dyads (*t*(19) = 3.68, *p* = 0.002). Consistent results were found in the ANOVA on RTs: significant effects of stimulus (*F*(2,38) = 15.90, *p* < 0.001, *η*_p_^2^ = 0.45) and orientation (*F*(1,19) = 24.13, *p* < 0.001, *η*_p_^2^ = 0.56) and a significant interaction (*F*(2,38) = 4.22, *p* = 0.022, *η*_*p*_^*2*^ = 0.18). In the RTs, the inversion effect was significant in all conditions (single bodies: *t*(19) = 4.60, *p* < 0.001; facing dyads, *t*(19) = 5.14, *p* < 0.001; nonfacing dyads, *t*(19) = 3.73, *p* = 0.001); but, again, it was significantly larger for facing dyads than for nonfacing dyads (*t*(19) = 2.72, *p* = 0.001).

**Figure 4.**
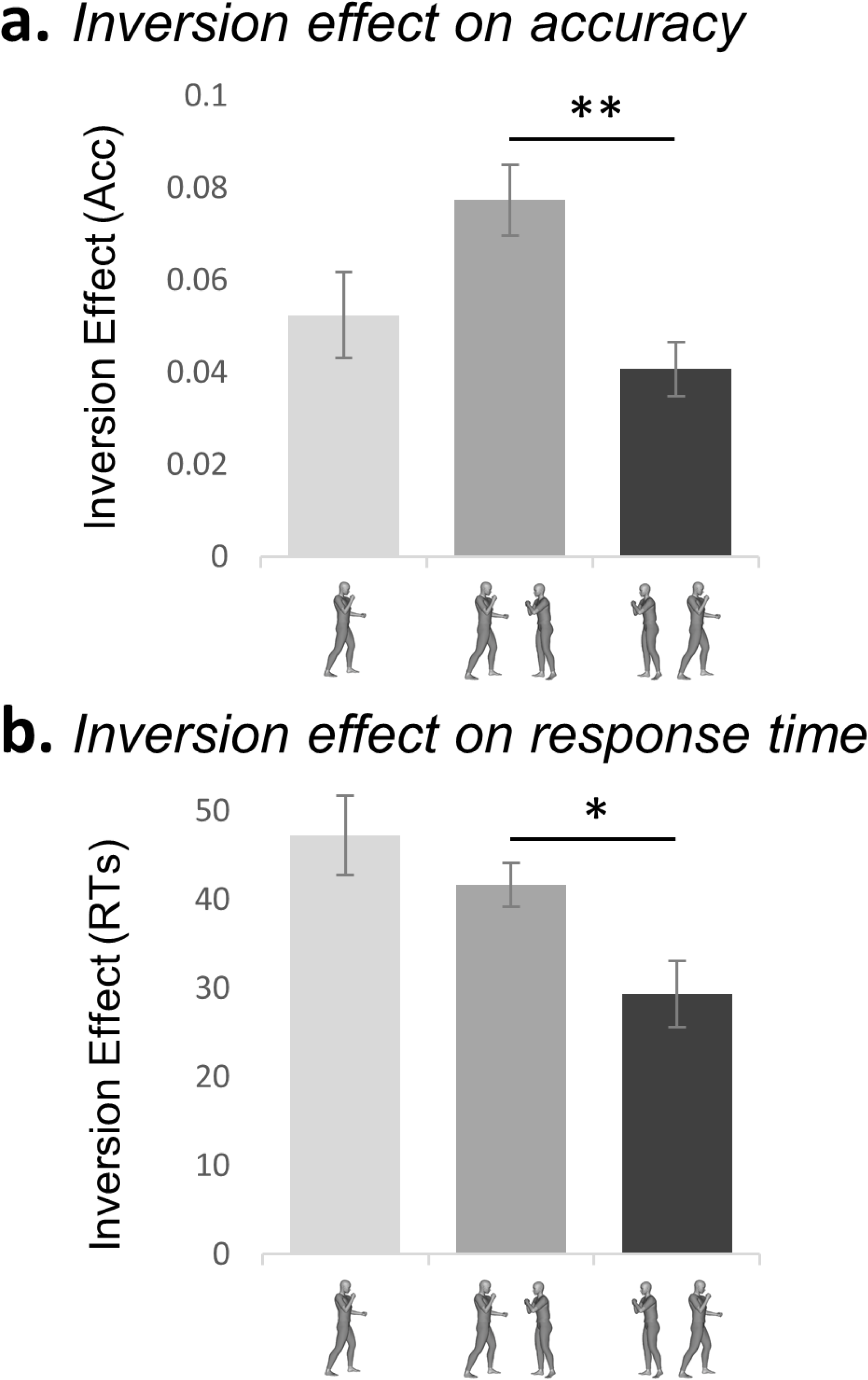
Inversion effect for facing and non-facing dyads. Inversion effect (proportion of correct responses with upright minus inverted trials ± within-subjects normalized *SEM*) for **(a)** accuracy (ACC) and **(b)** response time (RTs) as a function of the stimulus: single body, facing dyad and non-facing dyad. \**p* ≤ 0.05; \*\**p* ≤ 0.01.

Comparable results were obtained when considering only the 15 participants who took part in the fMRI experiment (accuracy: effect of stimulus, *F*(2,28) = 3.71, *p* = 0.037, *η*_*p*_^*2*^ = 0.21; effect of orientation: *F*(1,14) = 8.68, *p* = 0.011, *η*_*p*_^*2*^ = 0.38; interaction: *F*(2,28) = 3.58, *p* = 0.041, *η*_*p*_^*2*^ = 0.20; RTs: effect of stimulus, *F*(2,28) = 8.43, *p* = 0.001, *η*_*p*_^*2*^ = 0.38, effect of orientation, *F*(1,14) = 22.91, *p* < 0.001, *η*_*p*_^*2*^ = 0.62, interaction, *F*(2,38) = 3.15, *p* = 0.058, *η*_*p*_^*2*^ = 0.18).

The analysis on chair trials demonstrated that the effect of positioning on the cost of inversion (facing vs. nonfacing) was driven by the representation of the spatial relation between bodies, rather than merely their physical orientations. In fact, no difference in the inversion effect was found across different configurations of chair-stimuli (single chair, facing and nonfacing pairs). An ANOVA on accuracy values showed a main effect of stimulus (*F*(2,38) = 3.94, *p* = 0.028, *η*_*p*_^*2*^ = 0.17), but no effect of orientation (*F*(1,19) = 1.15, *p* > 0.250, *η*_*p*_^*2*^ = 0.06), or an interaction between them (*F*(2,38) = 2.47, *p* = 0.10, *η*_*p*_^*2*^ = 0.11). The ANOVA on RTs showed no effects of stimulus (*F*(2,38) = 2.75, *p* = 0.077, *η*_*p*_^*2*^ = 0.13), a significant effect of orientation (*F*(1,19) = 30.08, *p* < 0.001, *η*_p_ ^*2*^ = 0.61), but no interaction between the two factors (*F*(2,38) = 2.52, *p* = 0.094, *η*_*p*_^*2*^ = 0.12).

The increased inversion effect for facing dyads, as an index of efficiency in visual perception, may be the behavioral counterpart of the increased neural response for facing dyads in the body-selective cortex. Although the designs of the fMRI and the behavioral experiments were profoundly different (for one thing, only the latter involved inverted bodies), we tested the correlation between the behavioral and the neural effects in the 15 participants who took part to both the fMRI and the behavioral session. This analysis, across all the face/body-selective ROIs, yielded no significant effect (EBA: *mean rho* = −0.07, *t*(14) = −1.74 *p* = 0.105; FBA: *mean rho* = −0.07, *t*(14) = −1.69 *p* = 0.114, OFA: *mean rho* = −0.05, *t*(14) = −1.10, *p* = 0.292; and FFA: *mean rho* = −0.04, *t*(14) < 1, *n.s.*).

## Discussion

We investigated whether the relative positioning of two bodies –facing toward or away from each other–affected neural and behavioral signatures of body perception. Our findings demonstrate that the well-established sensitivity of human vision to single bodies extends to configurations of multiple bodies, whose face-to-face positioning may cue interaction.

Three fMRI data analyses and the behavioral results converged on demonstrating that cortical areas specialized for body perception capture spatial relations between bodies. Particularly in the EBA, a stronger response to facing *vs.* nonfacing dyads was found with a group-level whole-brain contrast, a voxel-by-voxel conjunction analysis in the LOC, and an analysis within the functionally defined ROIs, controlling for specific effects of limb information. Visual sensitivity to a stimulus, which accounts for neural activity in dedicated brain structures, should also be captured in visual perception behavior (DiCarlo et al., 2012). From this perspective, the larger inversion effect for facing (vs. nonfacing) dyads critically contributes to anchoring the above neural effects to visual sensitivity. Thus, the two-body inversion effect provides a fourth piece of evidence in favor of a tuning of human vision to configurations of multiple, seemingly interacting bodies.

The stronger neural response to facing dyads echoes the category-specific effects for bodies and faces in occipito-temporal regions (Kanwisher et al., 1997; Downing et al., 2001), as well as the effect of multiple objects appearing in functionally relevant relations (e.g., a hammer tilted toward a nail) *vs.* multiple unrelated objects, in the general object-selective LOC (Roberts and Humphreys, 2010; Kim and Biederman, 2011; Baeck et al., 2013). What specific process, triggered by facing dyads, accounts for the increased neural response in the EBA?

In visual areas, increased activity for related (*vs.* unrelated) items might signal an integrated representation that would alter the linear combination of responses to individual items (Kubilius et al., 2015). Using MVPA, we measured the extent to which the neural representation of dyads deviated from the mean response to single items (Baeck et al., 2013; Baldassano et al., 2017; Kaiser and Peelen, 2018). We found that representations of both facing and nonfacing dyads in the EBA matched the mean neural representation of single bodies. Ruling out the implementation of integrative processes, the processing of facing dyads in the EBA resembles the processing of single bodies, which trigger more activity than the scrambled set of their parts, and are linearly combined with the head-information in a whole-person representation (Kaiser et al., 2014; Brandman and Yovel, 2016; Harry et al., 2016). Tentatively, an indication of integrative processes for facing dyads was rather found in the prefrontal component of the network that responded to facing dyads more strongly than to nonfacing dyads. Here, the representation of dyads as a linear combination of single-body information, found for nonfacing dyads, was disrupted for facing dyads.

While we found no evidence of integrative processes in the EBA, we showed that the basic function of this area in encoding body shape or posture (Downing et al., 2006) was enhanced when a single body appeared in the context of another facing body. Discrimination of single bodies in facing dyads (but not in nonfacing dyads) was accurate, despite three detrimental factors. First, there was high visual similarity across bodies. Second, the instruction to fixate a cross and detect a color change diverted participants’ attention from bodies. Third, the neural patterns used for classification were obtained from blocks in which a given stimulus and its flipped versions were randomly interleaved. Thus, our analysis implemented a stringent test of body discrimination, which entailed the encoding of a given (unattended) posture across two different views. In this analysis, dyads could be successfully classified only if the single-body representation was particularly accurate. This was the case exclusively for the representation of single bodies in facing dyads in the EBA.

The representational sharpening in the EBA suggests an effect of visual context, whereby a body provides a meaningful context to another seemingly interacting body, enriching, or sharpening, its representation. The advantage of the neural representation of objects seen in a meaningful, or in their natural, context has been illustrated by visual context effects such as the word superiority effect (Reicher, 1969) and the face superiority effect (Homa et al., 1976). These effects demonstrate that an object (e.g., a nose) is identified better when seen in its regular *vs.* irregular context (i.e., a face). Furthermore, neuroimaging approaches have shown representational sharpening in the visual cortex for stimuli that meet subjective expectations, or an internal template, by appearing in the expected or regular context (Kok et al., 2012; Brandman and Peelen, 2017). Note that neural sharpening for single bodies in facing dyads is congruent with the lack of integrative processes in the EBA, as the latter effect implies *de facto* that the EBA preserves the information about single bodies intact: in the EBA, representation of single bodies in facing dyads is not only preserved but also enhanced.

Consistent with the fMRI results, the two-body inversion effect suggests that a facing dyad matches some sort of internal template that makes it easier to recognize, and particularly susceptible to spatial perturbation through inversion. The difference in the inversion effect between facing and nonfacing dyads is reminiscent of the difference between bodies and scrambled bodies (for which the body-inversion effect is reduced; Reed et al., 2003): like scrambled bodies, nonfacing dyads would break the expected configuration of two spatially close bodies, thus reducing the cost of inversion. The analogy also holds in light of similar neural effects for single bodies and facing dyads, such as the increased response relative to their scrambled/nonfacing counterpart and the linear combination of the component parts (body-parts/multiple bodies).

While both the neural and behavioral effects uncovered mechanisms specifically involved in processing of facing (vs. nonfacing) dyad configurations, we failed to find a correlation between neural response and inversion effect. Since inverted bodies were not included in the fMRI design, it is possible that the two measures captured qualitatively different aspects of neural processing. In any case, even in the more classic example of single bodies, the neural correlates of the inversion effect remain elusive: the body-inversion effect has been only inconsistently associated with the FFA, but not with the EBA (Brandman and Yovel, 2010; Brandman and Yovel, 2016).

In summary, our results showed that, in the EBA, facing dyads evoked an overall stronger response than nonfacing dyads, and sharpened the representation of single bodies. These findings prompt several questions for future research. First, it is important to test whether the current effects generalize to a large and more naturalistic set of body poses, beyond the eight used here. Second, the FFA and FBA registered the spatial relation between bodies with a stronger response to facing *vs.* nonfacing dyads, like the EBA, but showed no representational sharpening for single bodies in facing dyads or discrimination of individual body dyads. In the FFA, lack of discrimination of a single body or the dyad is not surprising given that all bodies had identical head/faces. However, the contribution of face/head *vs.* body, and the role of the different aspects of the face- and body-selective cortex in the effects of body positioning require further investigation. Moreover, the dissociation of the two effects (increased activation and representational sharpening) raises the possibility that different regions perform different computations, and/or that the two effects are signatures of different processes. Relatedly, we somehow implied that, in the EBA, representational sharpening might account for the increased response to facing dyads. However, an alternative interpretation is possible. Behavioral research suggests that spatial relations between bodies (facing *vs.* nonfacing) are captured early and automatically –i.e., before an elaborate, conscious recognition of actions from body postures (Papeo et al., 2017). Instead, sharpening of single objects’ representation has been characterized as the effect of top-down mechanisms on the visual cortex (Kok et al., 2012). Thus, in a hypothetical architecture, regions such as the EBA (and the FFA/FBA) could capture spatial relations between bodies, yielding stronger neural responses to facing dyads, and feed this information to higher-level systems (e.g., for action understanding). Higher-level systems could, in turn, put pressure on the encoding of single body postures, yielding representational sharpening in the EBA. Such a model may account for the recruitment of the prefrontal areas as well as the middle/superior temporal gyrus, which is an important territory for action understanding (Wurm and Lingnau, 2015; Wurm et al., 2017), with patches of selectivity for social interactions (Isik et al., 2017; Walbrin et al., 2018; Walbrin and Koldewyn, 2019).

Laying the foundations of future research, this report of specialized processes for a two-body shape demonstrates that the spatial relation of a body with another nearby body is captured in the same regions that serve body perception, and affects the way in which bodies are processed. It suggests that, beyond socially relevant entities, human vision is specialized for processing multi-body configurations, where the relative positioning of bodies signals the unfolding of a social event. Special sensitivity to socially relevant spatial relations between people contributes to characterizing human vision as *social*.

## Acknowledgments

The authors thank Jean-Remy Hochmann and Moritz Wurm for comments on an earlier version of this manuscript. This work was supported by a European Research Council Starting Grant awarded to L.P. (Project: THEMPO, Grant Agreement 758473).

## References

Adams RB, Ambady N, Nakayama K, Shimojo S (2010) The Science of Social Vision. Oxford University Press.

Aminoff E, Gronau N, Bar M (2007) The Parahippocampal Cortex Mediates Spatial and Nonspatial Associations. Cereb Cortex 17:1493–1503.

Ashburner J (2007) A fast diffeomorphic image registration algorithm. Neuroimage 38:95–113.

Baeck A, Wagemans J, Op de Beeck HP (2013) The distributed representation of random and meaningful object pairs in human occipitotemporal cortex: The weighted average as a general rule. Neuroimage 70:37–47.

Baldassano C, Beck DM, Fei-Fei L (2017) Human-Object Interactions Are More than the Sum of Their Parts. Cereb Cortex 27:2276–2288.

Baron-Cohen S, Baldwin DA, Crowson M (1997) Do Children with Autism Use the Speaker’s Direction of Gaze Strategy to Crack the Code of Language? Child Dev 68:48–57.

Birmingham E, Bischof WF, Kingstone A (2009) Saliency does not account for fixations to eyes within social scenes. Vision Res 49:2992–3000.

Bonatti LL, Frot E, Mehler J (2005) What Face Inversion Does to Infants’ Counting Abilities. Psychol Sci 16:506–510.

Brainard DH (1997) The Psychophysics Toolbox. Spat Vis 10:433–436.

Brandman T, Peelen M V. (2017) Interaction between Scene and Object Processing Revealed by Human fMRI and MEG Decoding. J Neurosci 37:7700–7710.

Brandman T, Yovel G (2010) The Body Inversion Effect Is Mediated by Face-Selective, Not Body-Selective, Mechanisms. 30:10534–10540.

Brandman T, Yovel G (2016) Bodies are Represented as Wholes Rather Than Their Sum of Parts in the Occipital-Temporal Cortex. Cereb Cortex 26:530–543.

Chang C-C, Lin C-J (2011) LIBSVM: a library for support vector machines. ACM Trans Intell Syst Technol 2:1–27.

Dale AM (1999) Optimal experimental design for event-related fMRI. Hum Brain Mapp 8:109–114.

Desikan RS, Ségonne F, Fischl B, Quinn BT, Dickerson BC, Blacker D, Buckner RL, Dale AM, Maguire RP, Hyman BT, Albert MS, Killiany RJ (2006) An automated labeling system for subdividing the human cerebral cortex on MRI scans into gyral based regions of interest. Neuroimage 31:968–980.

Diamond R, Carey S (1986) Why Faces Are and Are Not Special : An Effect of Expertise. J Exp Psychol Gen 115:107–117.

DiCarlo JJ, Zoccolan D, Rust NC (2012) How Does the Brain Solve Visual Object Recognition? Neuron 73:415–434.

Ding X, Gao Z, Shen M (2017) Two Equals One: Two Human Actions During Social Interaction Are Grouped as One Unit in Working Memory. Psychol Sci 28:1311–1320.

Downing PE, Bray D, Rogers J, Childs C (2004) Bodies capture attention when nothing is expected. Cognition 93:B27–B38.

Downing PE, Jiang Y, Shuman M, Kanwisher N (2001) A cortical area selective for visual processing of the human body. Science 293:2470–2473.

Downing PE, Peelen M V. (2011) The role of occipitotemporal body-selective regions in person perception. Cogn Neurosci 2:186–203.

Downing PE, Peelen M V., Wiggett AJ, Tew BD (2006) The role of the extrastriate body area in action perception. Soc Neurosci 1:52–62.

Faul F, Erdfelder E, Lang AG, Buchner A (2007) G*Power 3: A flexible statistical power analysis program for the social, behavioral, and biomedical sciences. Behav Res Methods 39:175–191.

Fischl B (2012) FreeSurfer. Neuroimage 62:774–781.

Friston KJ, Ashburner J, Kiebel S, Nichols T, Penny WD (2007) Statistical parametric mapping : the analysis of funtional brain images. Elsevier/Academic Press.

Friston KJ, Penny WD, Glaser DE (2005) Conjunction revisited. Neuroimage 25:661–667.

Friston KJ, Williams S, Howard R, Frackowiak RSJ, Turner R (1996) Movement-Related effects in fMRI time-series. Magn Reson Med 35:346–355.

Gauthier I, Skudlarski P, Gore JC, Anderson AW (2000) Expertise for cars and birds recruits brain areas involved in face recognition. Nat Neurosci 3:191–197.

Glanemann R, Zwitserlood P, Bölte J, Dobel C (2016) Rapid apprehension of the coherence of action scenes. Psychon Bull Rev 23:1566–1575.

Gobbini MI, Haxby J V. (2007) Neural systems for recognition of familiar faces. Neuropsychologia 45:32–41.

Gomez J, Natu V, Jeska B, Barnett M, Grill-Spector K (2018) Development differentially sculpts receptive fields across early and high-level human visual cortex. Nat Commun 9:788.

Graziano MSA, Kastner S (2011) Human consciousness and its relationship to social neuroscience: A novel hypothesis. Cogn Neurosci 2:98.

Hafri A, Papafragou A, Trueswell JC (2013) Getting the gist of events: Recognition of two-participant actions from brief displays. J Exp Psychol Gen 142:880–905.

Hafri A, Trueswell JC, Strickland B (2018) Encoding of event roles from visual scenes is rapid, spontaneous, and interacts with higher-level visual processing. Cognition 175:36–52.

Harry BB, Umla-runge K, Lawrence AD, Graham KS, Downing PE (2016) Evidence for Integrated Visual Face and Body Representations in the Anterior Temporal Lobes. J Cogn Neurosci 28:1178–1193.

Haxby J V., Gobbini MI, Furey ML, Ishai A, Schouten JL, Pietrini P (2001) Distributed and Overlapping Representations of Faces and Objects in Ventral Temporal Cortex. Science 293:2425–2430.

Homa D, Haver B, Schwartz T (1976) Perceptibility of schematic face stimuli: Evidence for a perceptual Gestalt. Mem Cognit 4:176–185.

Isik L, Koldewyn K, Beeler D, Kanwisher N (2017) Perceiving social interactions in the posterior superior temporal sulcus. Proc Natl Acad Sci 114:9145–9152.

Jenkinson M, Beckmann CF, Behrens TEJ, Woolrich MW, Smith SM (2012) FSL. Neuroimage 62:782–790.

Kaiser D, Peelen M V. (2018) Transformation from independent to integrative coding of multi-object arrangements in human visual cortex. Neuroimage 169:334–341.

Kaiser D, Stein T, Peelen M V. (2014) Object grouping based on real-world regularities facilitates perception by reducing competitive interactions in visual cortex. Proc Natl Acad Sci 111:11217–11222.

Kanwisher N, McDermott J, Chun MM (1997) The fusiform face area: a module in human extrastriate cortex specialized for face perception. J Neurosci 17:4302–4311.

Kim JG, Biederman I (2011) Where do objects become scenes? Cereb Cortex 21:1738–1746.

Kok P, Jehee JFM, de Lange FP (2012) Less Is More : Expectation Sharpens Representations in the Primary Visual Cortex. Neuron 75:265–270.

Kubilius J, Baeck A, Wagemans J, Op de Beeck HP (2015) Brain-decoding fMRI reveals how wholes relate to the sum of parts. Cortex 72:5–14.

Lee S, Kable JW (2018) Simple but robust improvement in multivoxel pattern classification. PLoS One 13:1–15.

Maurer D, Le Grand R, Mondloch CJ (2002) The many faces of configural processing. Trends Cogn Sci 6:255–260.

McCarthy P (2019) FSLeyes. Zenodo http://doi.org/10.5281/zenodo.1470761.

Natu VS, Barnett MA, Hartley J, Gomez J, Stigliani A, Grill-Spector K (2016) Development of Neural Sensitivity to Face Identity Correlates with Perceptual Discriminability. J Neurosci 36:10893–10907.

Neri P, Luu JY, Levi DM (2006) Meaningful interactions can enhance visual discrimination of human agents. Nat Neurosci 9:1186–1192.

New J, Cosmides L, Tooby J (2007) Category-specific attention for animals reflects ancestral priorities, not expertise. Proc Natl Acad Sci 104:16598–16603.

Oosterhof NN, Connolly AC, Haxby J V. (2016) CoSMoMVPA: Multi-Modal Multivariate Pattern Analysis of Neuroimaging Data in Matlab/GNU Octave. Front Neuroinform 10:1–27.

Papeo L, Abassi E (2019) Seeing social events: The visual specialization for dyadic human–human interactions. J Exp Psychol Hum Percept Perform 45:877–888.

Papeo L, Goupil N, Soto-Faraco S (2019) Visual Search for People Among People. Psychol Sci 30:1483–1496.

Papeo L, Stein T, Soto-Faraco S (2017) The Two-Body Inversion Effect. Psychol Sci 28:369–379.

Quadflieg S, Gentile F, Rossion B (2015) The neural basis of perceiving person interactions. Cortex 70:5–20.

Quadflieg S, Koldewyn K (2017) The neuroscience of people watching: How the human brain makes sense of other people’s encounters. Ann N Y Acad Sci 1396:166–182.

Reed CL, Stone VE, Bozova S, Tanaka J (2003) The body-inversion effect. Psychol Sci 14:302–308.

Reicher GM (1969) Perceptual recognition as a function of meaningfulness of stimulus material. J Exp Psychol 81:275–280.

Roberts KL, Humphreys GW (2010) Action relationships concatenate representations of separate objects in the ventral visual system. Neuroimage 52:1541–1548.

Saxe R, Brett M, Kanwisher N (2006) Divide and conquer: A defense of functional localizers. Neuroimage 30:1088–1096.

Schwarzlose RF, Swisher JD, Dang S, Kanwisher N (2008) The distribution of category and location information across object-selective regions in human visual cortex. Proc Natl Acad Sci 105:4447–4452.

Soares JM, Magalhães R, Moreira PS, Sousa A, Ganz E, Sampaio A, Alves V, Marques P, Sousa N (2016) A Hitchhiker’s Guide to Functional Magnetic Resonance Imaging. Front Neurosci 10:515.

Stigliani A, Weiner KS, Grill-Spector K (2015) Temporal Processing Capacity in High-Level Visual Cortex Is Domain Specific. J Neurosci 35:12412–12424.

Taylor JC, Wiggett AJ, Downing PE (2007) Functional MRI Analysis of Body and Body Part Representations in the Extrastriate and Fusiform Body Areas. J Neurophysiol 98:1626–1633.

Vestner T, Tipper SP, Hartley T, Over H, Rueschemeyer S-A (2019) Bound together: Social binding leads to faster processing, spatial distortion, and enhanced memory of interacting partners. J Exp Psychol Gen 148:1251–1268.

Walbrin J, Downing P, Koldewyn K (2018) Neural responses to visually observed social interactions. Neuropsychologia 112:31–39.

Walbrin J, Koldewyn K (2019) Dyadic interaction processing in the posterior temporal cortex. Neuroimage 198:296–302.

Wang L, Mruczek REB, Arcaro MJ, Kastner S (2015) Probabilistic Maps of Visual Topography in Human Cortex. Cereb Cortex 25:3911–3931.

Weiner KS, Barnett MA, Lorenz S, Caspers J, Stigliani A, Amunts K, Zilles K, Fischl B, Grill-Spector K (2017) The Cytoarchitecture of Domain-specific Regions in Human High-level Visual Cortex. Cereb Cortex 27:146–161.

Wurm MF, Caramazza A, Lingnau A (2017) Action Categories in Lateral Occipitotemporal Cortex Are Organized Along Sociality and Transitivity. J Neurosci 37:562–575.

Wurm MF, Lingnau A (2015) Decoding Actions at Different Levels of Abstraction. J Neurosci 35:7727–7735.

Yin (1969) Looking At Upside-Down Faces 1. J Exp Psychol 81:141–145.

